# TIMELINE OF BRAIN ALTERATIONS IN ALZHEIMER’S DISEASE ACROSS THE ENTIRE LIFESPAN

**DOI:** 10.1101/229195

**Authors:** Pierrick Coupé, José Vicente Manjón, Enrique Lanuza, Gwenaelle Catheline, for the Alzheimer’s Disease Neuroimaging Initiative

**Affiliations:** Univ. Bordeaux, LaBRI, UMR 5800, PICTURA, F-33400 Talence, France.; CNRS, LaBRI, UMR 5800, PICTURA, F-33400 Talence, France.; Instituto Universitario de Tecnologías de la Información y Comunicaciones (ITACA), Universitat Politècnica de València, Camino de Vera s/n, 46022 Valencia, Spain; Univ. Valencia, Dept. of Cell Biology, Burjassot 46100, Valencia, Spain; Univ. Bordeaux, CNRS, EPHE PSL Research Univ., INCIA, UMR 5283, F-33000 Bordeaux

## Abstract

Brain imaging studies have shown that progressive cerebral atrophy characterized the development of Alzheimer’s Disease (AD). The key question is how long before the diagnosis of AD the neurodegenerative process started leading to these structural alterations. To answer this question, we proposed an innovative way by inferring brain structure volume trajectories across the entire lifespan using massive number of MRI (N=4714). Our study provides evidences of early divergence of the AD model from the control model for the hippocampus before 40 years, followed by the lateral ventricles and the amygdala around 40 years for the AD model. Moreover, our lifespan investigation reveals the dynamic of the evolution of these biomarkers and suggest close abnormality trajectories for the hippocampus and the amygdala. Finally, our results highlight that the temporal lobe atrophy, a key biomarker in AD, is a very early pathophysiological event potentially associated to early life exposures to risk factors.

---

Alzheimer’s disease (AD) is the most prevalent form of dementia in persons older than 65 years [1]. Cognitive impairment, mainly related to memory deficits, is the most common manifestation of this disease [2]. Available neuroimaging evidence suggests that the neuropathological alterations underlying AD probably begin much earlier than the appearance of clinical symptoms and years before clinical diagnosis [3]. From these results, it appeared that the pharmacological management was finally implemented in patients with a largely advanced neurodegenerative process, making it difficult to fight against pathological progression. In this context, the concept of disease-modifying therapies is emerging and the search for early biomarkers of these alterations is currently a hot topic of research [4]. Neurodegeneration, assessed by the level of cerebral atrophy, is one of these biomarkers. In recent decades, several studies have investigated neurodegeneration in the prodromal phase of Alzheimer’s disease. However, studying the prodromal phase of the disease, which is an asymptomatic phase, is a difficult task. Indeed, this type of study is based on subjects with rare autosomal dominant mutations associated with a high risk of developing dementia [5–7] or on longitudinal studies with long follow-up in which brain imaging can be performed before the appearance of clinical symptoms (i.e., memory impairment) [8–10]. In these previous long follow-up studies, the starting point of the neurodegeneration was not determined since incident cases already present brain morphometric differences at baseline 7 or 10 years before the diagnosis [11, 12]. Therefore, the key question remains, how long before the AD brain trajectory diverges from the cognitively normal model? To answer this question, an alternative approach is to build an extrapolated lifespan model of AD brain structures by using large-scale databases. Indeed, epidemiological studies indicate that late dementia is associated with early exposure to risk factors at midlife, highlighting the need to consider brain biomarkers throughout the entire lifespan [13–15]. Therefore, we propose to take advantage of the new paradigm of BigData sharing in neuroimaging [16] to analyze publically available large-scale databases containing subjects from a wide age range covering the entire lifespan.

Recently, we used such BigData approach to propose an analysis of brain trajectory across the entire lifespan using N=2944 MRI of cognitively normal subjects (CN) [17]. Herein, we present a study following a similar approach to perform lifespan analysis of the timeline of brain atrophy in AD. To this end, we propose to build an extrapolated model of AD for brain structures. We assume that the neurodegenerative process is slow and progressive. Consequently, to build our lifespan AD model we used N=3262 MRI data, 1385 from AD and Mild Cognitive Impairment (MCI) patients (from 55y to 96y) and 1877 from healthy/asymptomatic subjects younger than them (from 9 months to 55y). The proposed approach can be viewed as a conservative lifespan model of AD since CN are used as young asymptomatic AD subjects, in agreement with studies suggesting brain alterations at presymptomatic stage several years before diagnosis or MCI stage [11, 12]. We have focused the present study on temporal lobe structures such as the hippocampus and amygdala – known to be affected in AD [18] – and the lateral ventricles, also a known AD biomarker [19, 20]. We also included global white matter and gray matter and subcortical structures – thalamus, accumbens, caudate, putamen and globus pallidus.

## Results

Figure 1 presents trajectories of all considered structures for AD/MCI and CN groups (see online method for groups definition). This figure shows that hippocampus (HPC) and amygdala (AG) models present marked divergences between AD/MCI and CN, but also indicates that this divergence increases with age. Moreover, the divergence of control and pathological models for these structures occurs early around 40-45y. Lateral ventricles (LV) also exhibits early divergence – starting around 42y –between both models, however the distance between models decreases at advanced ages. Similarly, the thalamus presents an early but weak divergence that decreases at advanced ages. Pathological models of caudate and accumbens nuclei exhibit accelerated volume decreases from 50-60y. However, confidence intervals for these structures overlap again after 85y (see Table 1). For white matter (WM) and grey matter GM, AD/MCI models present an early accelerated aging compared to CN models around 45y. However, after 80y, CN models of brain tissues show an accelerated volume decreases. Consequently, confidence intervals of pathological and normal models overlap after 85 years (see Table 1). Finally, normal and pathological models for globus pallidus and putamen present similar trends.

**Figure 1.**
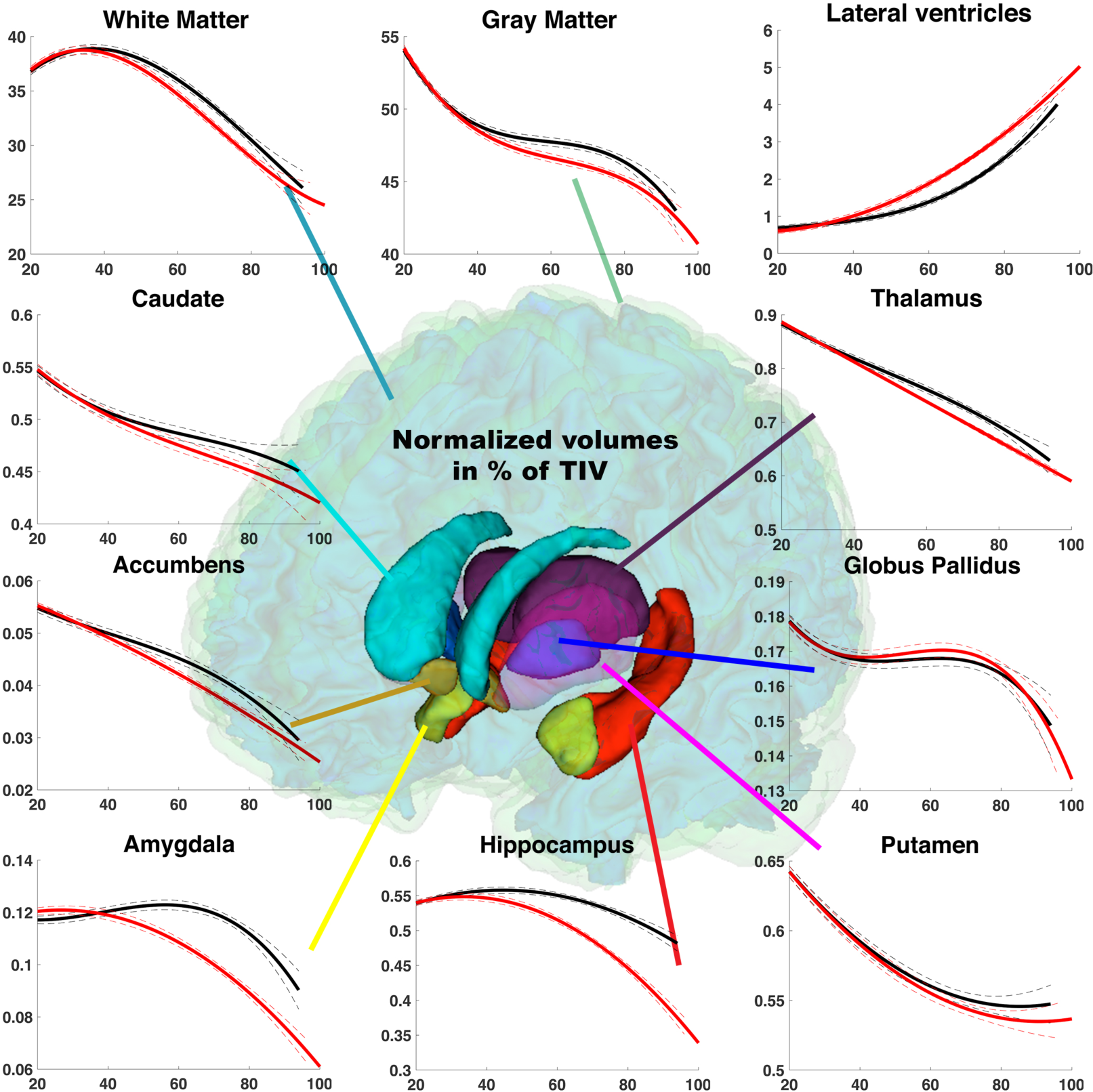
Trajectories based on relative volumes (% total intracranial volume) for brain cortical and subcortical structures across the entire lifespan. These volume trajectories are estimated according to the age of subjects. Model for CN group (N=2944) is in black and model for AD/MCI group (N=3262) is in red. The prediction bounds of the models are estimated with a confidence level at 95%.

**Table 1:**
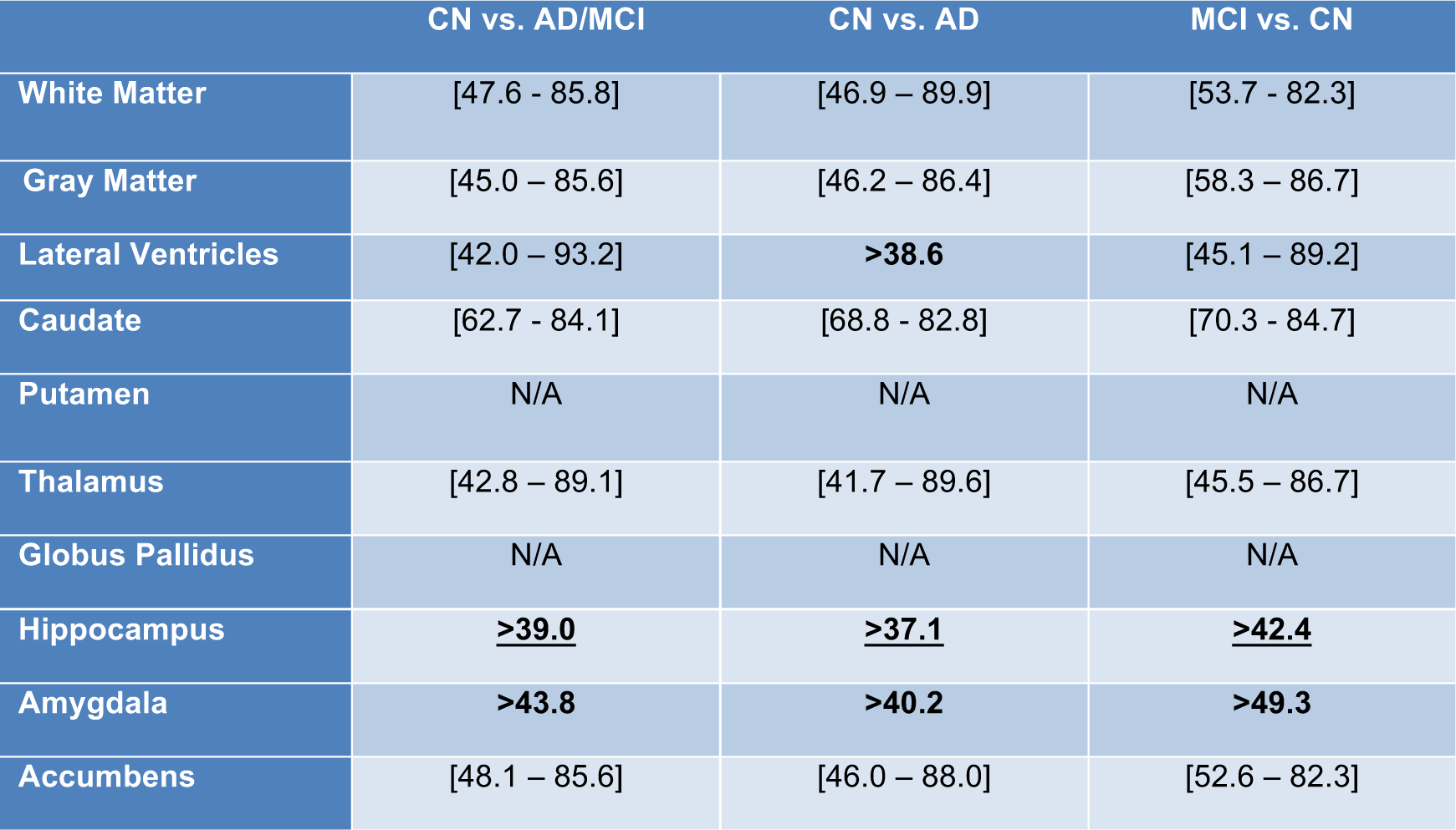
*Age range in years where confidence intervals of the predicted pathological models do not overlap with the predicted control models. The prediction bounds are estimated with a confidence level at 95%. Three model comparisons are presented CN (N=2944) vs. AD/MCI (N=3262), CN (N=2944) vs. AD (N=2303) and CN (N=2944) vs. MCI (N=2836)*.

Table 1 shows the age ranges where the confidence interval of the predicted pathological models (i.e., AD, MCI and AD/MCI) do not overlap with the confidence interval of the control models.

First, only HPC and AG trajectories present non-overlapping confidence intervals after trajectory divergence for all the studied pathological modes (i.e., AD/MCI, AD and MCI) (see Table 1). This is also valid for lateral ventricle but only when using the AD group. For all other considered structures, predicted confidence intervals overlap again at advanced ages around 80-90y.

Second, HPC is the first impacted deep gray structure with a trajectory divergence at 39y when using the AD/MCI group, 37y when using the AD group and 42y for the MCI group. The second structure impacted is the LV with a divergence point at 42y when using AD/MCI group, 39y for AD group and 45y for MCI group. Afterwards, thalamus (TH) trajectory divergence from control at 43y when using AD/MCI group, 42y for AD group and 45y for MCI group. AG trajectory divergence occurs then at 44y when using AD/MCI group, 41y when using AD group and 49y for MCI group. Impact on global GM and WM volume is observed later, with trajectories diverging at 45y and 48y respectively for the AD/MCI group, at 46y and 47y respectively for the AD group and at 58y and 54y respectively for the MCI group. Finally, accumbens and caudate trajectories diverge slightly later, but in a similar age range. Putamen and globus pallidus are the only deep gray matter structures for which trajectories do not diverge from CN across the entire lifespan.

To further analyze trajectories of well-known AD biomarkers, we propose an additional analysis focusing on the HPC, the LV and the AG. Figure 2 presents the trajectories of these structures for CN, AD and MCI groups. Moreover, relative rate of change and abnormality percentages are provided.

**Figure 2:**
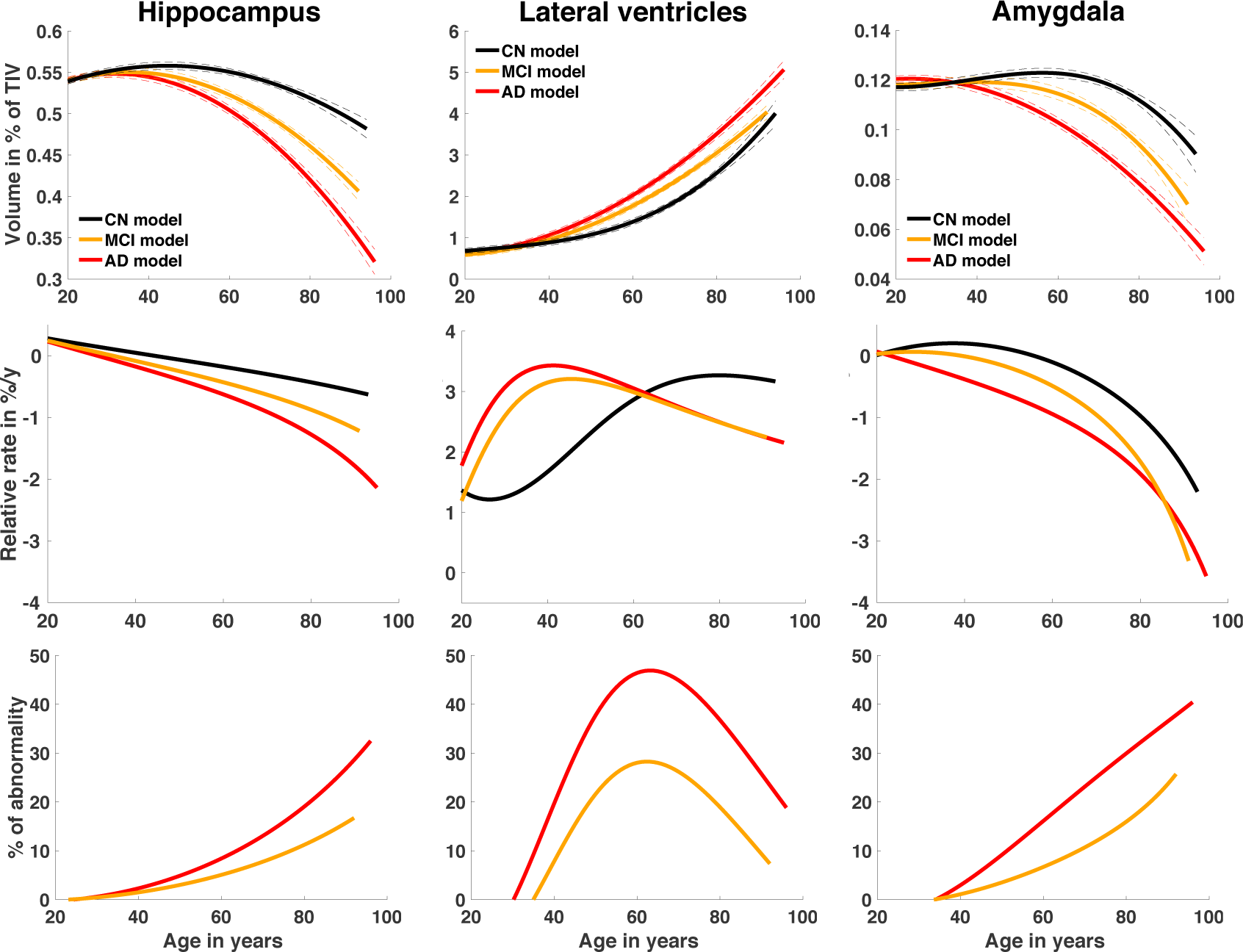
Hippocampus, lateral ventricles and amygdala trajectories for CN, AD and MCI models. The relative volumes (% total intracranial volume) are displayed according to the age in years across the entire lifespan. The prediction bounds are estimated with a confidence level at 95%. Relative rate of change is based on the first derivative of the model divided by the model and provided in % per year. Finally, percentage of abnormality is estimated as the difference between CN model and AD or MCI models divided by CN model. The model for CN group (N=2944) is displayed in black, the model for MCI group (N=2836) is displayed in yellow and the model for AD group (N=2303) is displayed in red.

First, divergence points for the AD models compared to the CN models are earlier than for the MCI models (see Table 1 for exact time). As expected, MCI trajectories are in between AD and CN ones. Second, when using relative rate of change, AG and LV exhibit a more pronounced relative change compared to HPC. The maximum relative changes for AD models of these structures are −3.6%/y for AG at 96y, −2.1%/y for HPC at 96y and 3.4%/y at 42y for LV. Contrary to HPC and AG, which show an increasing relative change with age, LV exhibits enlargement following an inverted U-shape. When considering abnormality percentage, an earlier abnormality increase is observed for HPC than for LV and AG. This abnormality reaches a maximum of 32% for the AD model at 96y. Abnormality appears later in life for LV and AG and follows very different patterns for both. The LV abnormality trajectory follows an inverted U-shape with a maximum of 47% at 63y for the AD model. The AG abnormality trajectory has similar trend to that of the HPC abnormality. AG reaches 40% of abnormality at 96y for the AD model. Therefore, while HPC abnormality starts first, AG presents a greater abnormality at advanced age. Moreover, the abnormality observed in LV is also important but its maximum is reached at 65y. Afterwards, percentage of abnormality of LV decreases to end at 19% at 96y for the AD model. Therefore, at late age, the LV shows lower abnormalities than those of the HPC and the AG at the same ages.

## Discussion

Our lifespan analysis based on large-scale datasets using inferred timeline of brain atrophy in AD indicates that the HPC is the first brain structure to exhibits a significant volume difference between cognitively normal subjects and subjects who will present clinical symptoms. This difference is detectable early in life, at 39y for the AD/MCI model and at 37y for the AD model. The HPC is followed by another temporal lobe region, the AG, which is different between the two groups at 44y for the AD/MCI model and at 40y for the AD model. It is noticeable that AG is undergoing larger changes proportionally to its size compared to HPC. Finally, the LV presents an early enlargement at 42y for the AD/MCI model and at 39y for the AD model. However, LV enlargement occurring during normal aging reduces the abnormality of this structure after 65y. Finally, TH shows early divergence at 43y for the AD/MCI model and at 42y for the AD model.

Our results presenting the HPC as the first brain region diverging in the preclinical stage of AD is in accordance with previous morphometric studies focused on the prodromal phase of the disease [9–12]. It is also in accordance with histopathological studies showing the temporal lobe as the starting point of the neurodegenerative process in AD [21]. In the long follow-up studies mentioned previously, authors observed that incident cases of AD present morphometric difference in the hippocampus at least 10 years before the diagnosis. In our study, the youngest subjects presenting clinical symptoms included in the AD model are 55 years old, while the pathological trajectories diverge from normal model 18 years and 16 years for the AD model and the AD/MCI model, respectively. This result suggests that the neurodegeneration of the hippocampus is present several years before the onset of cognitive deficit. Therefore, our model seems to confirm the presence of a long-lasting period of silence before the diagnosis of AD, as discussed in [22]. Moreover, our model indicates that the age of 40y is a critical period in the onset of the temporal lobe atrophy. Exposure to risk factors (such as diabetes and smoking) occurring at this lifetime period should be considered in future studies to evaluate their implication in the atrophy process [14]. It worth to note that all the results about HCP in this paper have been obtained using the EADC-ADNI harmonized protocol. Therefore, this study, to the best of our knowledge, is the largest analysis using this protocol to date.

The second temporal lobe region diverging from the cognitively normal subjects according to our model is the AG, which is different from CN at 40y for the AD model and at 44y for the MCI/AD models. Atrophy of this structure has been repeatedly described in AD subjects, with a rate of change less important [23] or similar [18] than hippocampal one. Notably, in the transgenic mouse model APPswe/ PS1dE9 of AD, the neurodegeneration in the amygdala even precedes that found in the hippocampus [24]. In our study, we found that the relative rate of change and abnormality were greater for AG than for the HPC at advanced ages. The early atrophy of the AG in the prodromal phase of AD is not surprising when considering the implication of emotion in memory. Indeed, the activity of basolateral and lateral nuclei of the AG is associated to a facilitation during the encoding phase and to an enhanced retrieval, these effects being mediated through the important interconnections between these structures and the HPC [25]. In addition, degradation of emotion processing ability is also observed in AD patients, as expected given the amygdala atrophy [26]. Moreover, the atrophy of the AG is likely contributing to the olfactory deficits associated with AD, since the cortical nuclei of the AG are associated with the processing of olfactory stimuli [27]. Hyposmia has been described in AD [28], and olfactory deficits can substantially precede cognitive symptoms [29]. However, it has to be taken into account that pathological alterations in AD occur also in other olfactory structures [30]. Finally, timeline atrophy of the HPC and the AG never overlaps across lifespan between the AD and CN models, in contrast to other deep gray matter structures investigated in this study. This result highlights the specificity all along life of the medial temporal lobe atrophy associated to the mnesic symptoms, which characterize the disease.

According to our results, the volume of the LV is also an early biomarker of AD, since its trajectory diverges at 39y for the AD model and at 42y for the AD/MCI model. The potential of LV volume as AD biomarkers has been previously mentioned over restricted periods [20]. In this study, by analyzing the change of LV abnormality over time, we showed that LV abnormality decreases after 65y. Therefore, the use of this biomarker is difficult for the late onset cases due to important LV enlargement occurring during normal aging. However, it may be useful to discriminate cases around 65y, an early age at which the AD diagnosis is particularly relevant because intervention is more effective in the early phases of the disease [31]. The importance of taking into account volume increase at advanced age in normal aging has been previously mentioned [19]. In this study, early divergence has been also observed for TH around 42y. Such thalamic atrophy was previously reported in AD literature [32]. However, we found that TH abnormality was very small (4.6% at 81y). This may explain why only a small number of studies have mentioned that this structure could be affected by AD because a large number of subjects are needed to detect such subtle atrophy. Consequently, TH does not appear to be an optimal biomarker for AD.

In this study, we proposed to process a massive number of cross-sectional MRI to investigate timeline atrophy of AD across the entire lifespan. The use of crosssectional data to analyze a dynamic process may appear not optimal. However, previous studies demonstrated that cross-sectional and longitudinal approaches produce similar age-related patterns in normal aging and a similar atrophy model in AD [33]. Compared to previous longitudinal investigations, our results are consistent with most previous findings on the importance and timing of atrophy in the HPC and the AG and the enlargement of the LV [20, 34]. Moreover, the obtained values for relative rates of change fit well within the expected range of annual atrophy rate reported in the longitudinal literature for both AD and CN [34–36]. With respect to the estimated time of divergence between the CN and AD models, there is no longitudinal or cross-sectional literature over the lifetime with which to compare. However, long follow-up population-based studies tracking cognition decline at preclinical stage provides evidences of a very long silent phase. This preclinical period was estimated from few years up to several decades before AD diagnosis that is in consistent with our results [37–39].

## Online Method

### Groups definition

This study aims at comparing normal and pathological trajectories of brain atrophy during AD progression across the entire lifespan to study the timeline. To this end, models were estimated on four different groups to generate CN, AD/MCI, AD and MCI trajectories.

- For CN trajectories, we used the N=2944 subjects from 9 months to 94y of the cognitively normal dataset as done in [17].
- For the AD/MCI trajectories, we used N=3262 samples. We mixed AD patients, with MCI patients considered being at an early stage of AD, and with young CN considered as presymptomatic subjects. We used 426 AD patients (from 55y to 96y), 959 MCI patients (from 55y to 92y) of the AD/MCI dataset and all the CN younger than 55y (i.e., 1877 samples). These subjects are included in the CN used for CN trajectory.
- For the AD trajectories, we used N=2303 samples. We mixed AD patients with young CN. More precisely, we used 426 AD patients (from 55y to 96y) of the AD/MCI datasets and all the CN younger than 55y (i.e., 1877 samples).
- For the MCI trajectories, we used N=2836 samples. Here, 959 MCI patients (from 55y to 92y) of the AD/MCI datasets were mixed with all the CN younger than 55y (i.e., 1877 samples).

### Datasets description

To study structures trajectory across the entire lifespan for CN and AD, we aggregated several open access databases to construct two datasets. In the following, CN and AD/MCI datasets will be described. The four previously described groups are built on these datasets.

#### Cognitively normal dataset (N=2944)

The cognitively normal dataset is composed of the 3296 T1-weight (T1w) MRI used in [17]. The composition of the nine open access databases used to build the control dataset is provided in Table 2. As explained in [17], after a demanding 3-stage quality control, only 2944 MRI were kept. The female proportion is 47% for the remaining subjects and the age range is [0.75 – 94] years.

**Table 2:** *Dataset description used for the CN models. This table provides the name of the dataset, the MR acquisition configuration, the number of considered image before and after QC, the gender proportion after QC and the average mean, standard deviation in parentheses and the interval in brackets*.

| DATASET | Before QC | After QC | Gender after QC | Age in years after QC |
| --- | --- | --- | --- | --- |
| **C-MIND** | 266 | 236 | F = 129<br>M = 107 | 8.44 (4.35)<br>[0.74-18.86] |
| **NDAR** | 612 | 382 | F = 174<br>M = 208 | 12.39 (5.94)<br>[1.08-49.92] |
| **ABIDE** | 528 | 492 | F = 84<br>M = 408 | 17.53 (7.83)<br>[6.50-52.20] |
| **ICBM** | 308 | 294 | F = 142<br>M = 152 | 33.75 (14.32)<br>[18-80] |
| **IXI** | 588 | 573 | F = 321<br>M = 252 | 49.52 (16.70)<br>[20.0- 86.2] |
| **OASIS** | 315 | 298 | F = 187<br>M = 111 | 45.34 (23.82)<br>[18 - 94] |
| **AIBL** | 236 | 233 | F = 121<br>M = 112 | 72.24 (6.73)<br>[60 - 89] |
| **ADNI 1** | 228 | 223 | F = 108<br>M = 115 | 75.96 (5.03)<br>[60 – 90] |
| **ADNI 2** | 215 | 213 | F = 113<br>M = 100 | 74.16 (6.39)<br>[56.3 - 89] |
| **Total** | **3296** | **2944** | **F = 1379 (47%)**<br>**M = 1565 (53%)** | **39.65 (26.62)**<br>**[0.74 - 94]** |

#### AD/MCI dataset (N=1385)

The AD/MCI dataset is composed of 426 AD patients and 959 MCI patients extracted from OASIS, AIBL, ADNI1 and ADNI2 databases. Details on clinical criterion for groups definition are provided in [40] for ADNI1, ADNI2 and AIBL and in [41] for OASIS. The composition of the databases used to build the AD/MCI dataset is provided in Table 3. After our 3-stage quality control, only 1385 MRI were kept. The female proportion is 44% for the remaining subjects and the age range is [55 – 96] years.

**Table 3:**
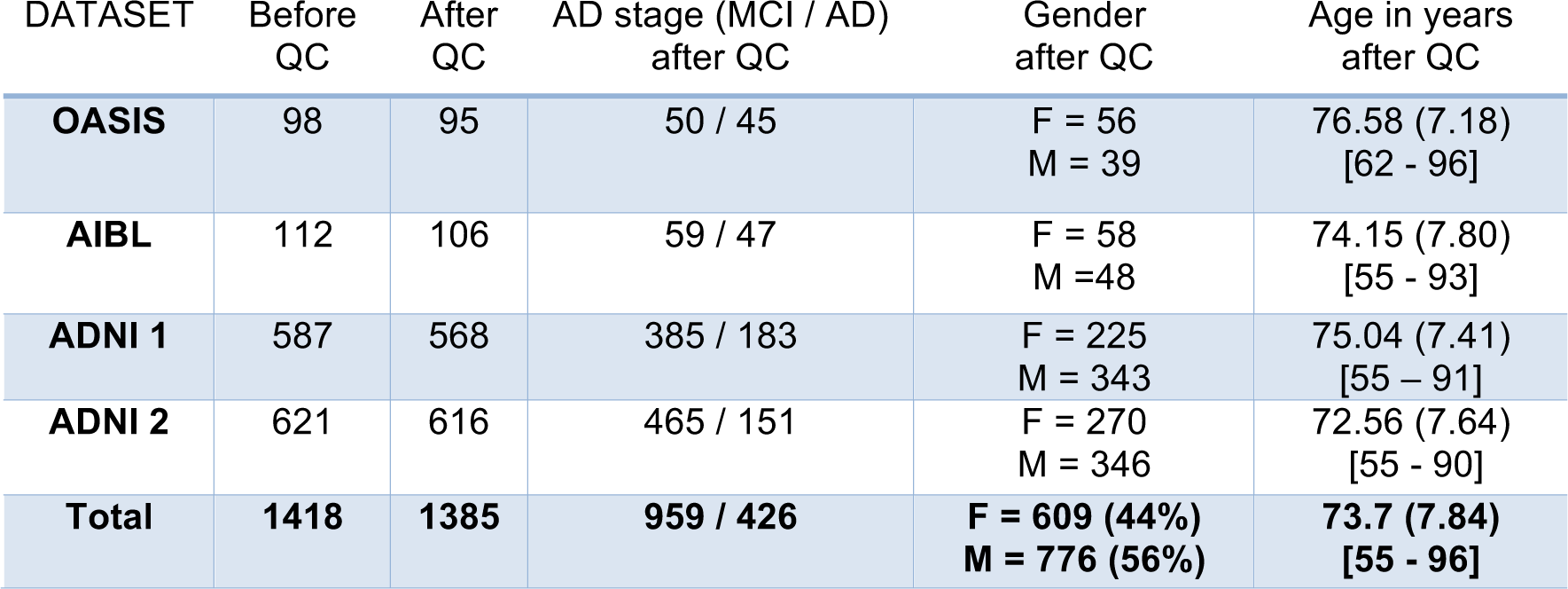
*Dataset description used for the AD models. This table provides the name of the dataset, the MR acquisition configuration, the number of considered image before and after QC, the gender proportion after QC and the average mean, standard deviation in parentheses and the interval in brackets*.

| DATASET | Before QC | After QC | AD stage (MCI / AD) after QC | Gender after QC | Age in years after QC |
| --- | --- | --- | --- | --- | --- |
| **OASIS** | 98 | 95 | 50 / 45 | F = 56<br>M = 39 | 76.58 (7.18)<br>[62 - 96] |
| **AIBL** | 112 | 106 | 59 / 47 | F = 58<br>M = 48 | 74.15 (7.80)<br>[55 - 93] |
| **ADNI 1** | 587 | 568 | 385 / 183 | F = 225<br>M = 343 | 75.04 (7.41)<br>[55 - 91] |
| **ADNI 2** | 621 | 616 | 465 / 151 | F = 270<br>M = 346 | 72.56 (7.64)<br>[55 - 90] |
| **Total** | **1418** | **1385** | **959 / 426** | **F = 609 (44%)**<br>**M = 776 (56%)** | **73.7 (7.84)**<br>**[55 - 96]** |

In the following, more details are provided about acquisition protocols of the different datasets used in this study.

- **C-MIND:** 266 images of control subjects from the C-MIND dataset (https://research.cchmc.org/c-mind/) are used in this study. All the 3D T1-weight (T1w) MPRAGE high-resolution MRI were acquired at the same site on a 3T scanner with spatial resolution of 1 mm^3^ acquired using a 32 channel SENSE head-coil.
- **NDAR**: 415 of control subjects from the Database for Autism Research (NDAR) (https://ndar.nih.gov) are used in this study. The T1w 3D MRI were acquired on 1.5T MRI and 3T scanners. In our experiments, we used the NIHPD (http://www.bic.mni.mcgill.ca/nihpd/info/dataaccess.html) dataset and 197 images of control subjects from the Lab Study 19 of National Database for Autism Research. For the NIHPD dataset, the 3D T1w SPGR MRI were acquired at six different sites with 1.5 Tesla systems by General Electric (GE) and Siemens Medical Systems with spatial resolution of 1 mm^3^. The 3D T1w MPRAGE MRI from the Lab Study 19 were scanned using a 3T Siemens Tim Trio scanner at each site with spatial resolution of 1 mm^3^
- **ABIDE**: 528 control subjects from the Autism Brain Imaging Data Exchange (ABIDE) dataset (http://fcon1000.projects.nitrc.org/indi/abide/) are used in this study. The MRI are T1w MPRAGE acquired at 20 different sites on 3T image and the details of acquisition, informed consent, and site-specific protocols are available on the website.
- **ICBM**: 308 normal subjects from the International Consortium for Brain Mapping (ICBM) dataset (http://www.loni.usc.edu/ICBM/) obtained through the LONI website are used in this study. The T1w MPRAGE MRI were acquired on a 1.5T Philips GyroScan imaging system (Philips Medical Systems, Best, The Netherlands) with spatial resolution of 1 mm^3^.
- **IXI**: 588 normal control from Information extraction from Images (IXI) database (http://brain-development.org/ixi-dataset/) are used in this study. The MRI are T1w images collected at 3 sites with 1.5 and 3T scanners with spatial resolution close to 1mm^3^.
- **OASIS**: 315 control subjects and 98 AD/MCI patients from the Open Access Series of Imaging Studies (OASIS) database (http://www.oasis-brains.org) are used in this study. The MRI are T1w MPRAGE image acquired on a 1.5T Vision scanner (Siemens, Erlangen, Germany) and resliced at 1mm^3^.
- **ADNI1**: 228 control subjects and 587 AD/MCI patients from the Alzheimer’s Disease Neuroimaging Initiative (ADNI) database (http://adni.loni.usc.edu) phase 1 are used in this study. These baseline MRI are T1w MPRAGE acquired on 1.5T scanners at 60 different sites across the United States and Canada with reconstructed spatial resolution of 1 mm^3^.
- **ADNI2**: 215 control subjects and 621 AD/MCI patients from the ADNI2 database (second phase of the ADNI project) are used in this study. The baseline MRI are T1w MPRAGE acquired on 3T Mr scanners with the standardized ADNI-2 protocol (www.loni.usc.edu) with spatial resolution close to 1mm^3^.
- **AIBL**: 236 control subjects and 112 AD/MCI patients from the Australian Imaging, Biomarkers and Lifestyle (AIBL) database (http://www.aibl.csiro.au/) are used in this study. The baseline MRI are T1w image acquired on 3T MR scanners with the ADNI protocol (http://adni.loni.ucla.edu/research/protocols/mri-protocols) and with custom MPRAGE sequence on the 1.5T scanners.

### Image processing

All the considered images were processed with the volBrain pipeline [42] (http://volbrain.upv.es). The volBrain system is a web-based online tool providing automatic brain segmentation and generating report summarizing the volumetric results. The full processing time is around 10 minutes. In the past 2 years, volBrain has processed online around 55.000 brains for approximately 1500 users. In a recent work, we compared volBrain pipeline with two well-known tools used on MR brain analysis (FSL and Freesurfer). We showed significant improvements in terms of both accuracy and reproducibility for intra and inter-scanner scan-rescan acquisitions [42]. The volBrain processing pipeline includes several steps to improve the quality of the input MR images and to homogenize their contrast and intensity range. The volBrain pipeline achieves the following preprocessing steps: 1) denoising using spatially adaptive non-local means [43], 2) rough inhomogeneity correction using N4 method [44], 3) affine registration to MNI152 space using ANTS software [45], 4) SPM based fine inhomogeneity correction [46] and 5) tissue based intensity standardization [47]. After preprocessing, the brain is segmented into several structures at different scales. First, the total intracranial volume (TIV) is obtained with NICE method [48]. Then, tissue classification is performed using the TMS method [47] and finally subcortical structures are estimated using the non-local label fusion method [49]. All the segmentation methods of volBrain are based on a library of 50 experts manually labelled cases (covering almost the entire lifespan). It is worth to note that the used manual hippocampus labeling followed the EADC-ADNI harmonized protocol which is the current consensus protocol for hippocampus segmentation in AD [50]. More details about volBrain pipeline can be found in [42]. Finally, a multi-stage quality control (QC) procedure was performed to carefully select subjects included. First, a visual assessment was done for all input images by checking screen shots of one sagittal, one coronal and one axial slice in middle of the 3D volume. Then, a visual assessment of processing quality was carried out by using the volBrain report which provides screenshots for each step of the pipeline. Finally, a last control was performed by individually checking with a 3D viewer all outliers detected using the estimated model (see [17] for more details).

### Statistical Analysis

Different model types were considered to estimate the final model of each structure. The candidate models were tested from the simplest to the most complex. A model type was kept as a potential candidate only when simultaneously F-statistic based on ANOVA (i.e., model vs. constant model) was significant (p<0.05) and when all its coefficients were significant using t-statistic (p<0.05). As in [17], the following model types were used as potential candidates:

1. Linear model

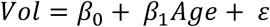
2. Quadratic model

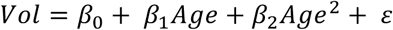
3. Cubic model

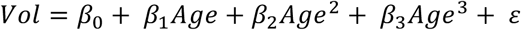
4. Linear hybrid model: exponential cumulative distribution for growth with linear model for aging

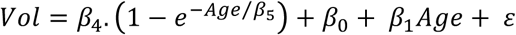
5. Quadratic hybrid model: exponential cumulative distribution for growth with quadratic model for aging

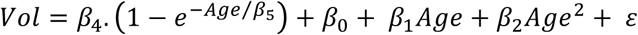
6. Cubic hybrid model: exponential cumulative distribution for growth with cubic model for aging

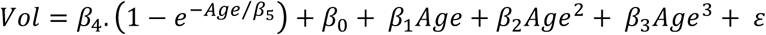 To select the best model type, we used the Bayesian Information Criterion (BIC) among kept candidate models – p<0.05 for ANOVA of the model vs. constant model and p<0.05 for T-test of all the coefficients. The BIC is a measure providing a trade-off between bias and variance to select the model explaining most of the data with a minimum number of parameters. Moreover, to compensate for variability introduced by head size difference, models were estimated on normalized volume in % of total intracranial volume (TIV). Left and right volumes were added to obtain the final volume structure. The prediction bounds were estimated with a confidence level at 95%. This model selection procedure was applied to all the considered structures. In this study, we studied the following brain structures: lateral ventricles (LV), caudate, thalamus (TH), accumbens, globus pallidus, amygdala (AG), hippocampus (HPC) and putamen. Moreover, tissue classification was used to obtain the global volume of white matter (WM) and gray matter (GM). All statistical tests were performed with Matlab© using default parameters. Afterwards, percentage of relative rate of change per year and percentage of abnormality were computed on the estimated models. The relative rate of change in percentage per year was computed as the first derivative of the model divided by the model and the abnormality as the difference
between pathological models and control model divided by control model.

## Acknowledgements

We gratefully acknowledge financial support from the French State, managed by the French National Research Ageny in the frame of the Investments for the future Program IdEx Bordeaux (ANR-10-IDEX-03-02, HL-MRI Project), Cluster of excellence CPU and TRAIL (HR-DTI ANR-10-LABX-57). We also want to think the financial support from the CNRS thanks to the multidisciplinary project Défi imag′In and the dedicated volBrain support.

Moreover, this work is based on multiple samples. We wish to thank all investigators of these projects who collected these datasets and made them freely accessible.

The C-MIND data used in the preparation of this article were obtained from the C-MIND Data Repository (accessed in Feb 2015) created by the C-MIND study of Normal Brain Development. This is a multisite, longitudinal study of typically developing children from ages newborn through young adulthood conducted by Cincinnati Children’s Hospital Medical Center and UCLA and supported by the National Institute of Child Health and Human Development (Contract #s HHSN275200900018C). A listing of the participating sites and a complete listing of the study investigators can be found at https://research.cchmc.org/c-mind.

The NDAR data used in the preparation of this manuscript were obtained from the NIH-supported National Database for Autism Research (NDAR). NDAR is a collaborative informatics system created by the National Institutes of Health to provide a national resource to support and accelerate research in autism. The NDAR dataset includes data from the NIH Pediatric MRI Data Repository created by the NIH MRI Study of Normal Brain Development. This is a multisite, longitudinal study of typically developing children from ages newborn through young adulthood conducted by the Brain Development Cooperative Group and supported by the National Institute of Child Health and Human Development, the National Institute on Drug Abuse, the National Institute of Mental Health, and the National Institute of Neurological Disorders and Stroke (Contract #s N01-HD02-3343, N01-MH9-0002, and N01-NS-9-2314, -2315, -2316, -2317, -2319 and -2320). A listing of the participating sites and a complete listing of the study investigators can be found at http://pediatricmri.nih.gov/nihpd/info/participating_centers.html.

The ADNI data used in the preparation of this manuscript were obtained from the Alzheimer’s Disease Neuroimaging Initiative (ADNI) (National Institutes of Health Grant U01 AG024904). The ADNI is funded by the National Institute on Aging and the National Institute of Biomedical Imaging and Bioengineering and through generous contributions from the following: Abbott, AstraZeneca AB, Bayer Schering Pharma AG, Bristol-Myers Squibb, Eisai Global Clinical Development, Elan Corporation, Genentech, GE Healthcare, GlaxoSmithKline, Innogenetics NV, Johnson & Johnson, Eli Lilly and Co., Medpace, Inc., Merck and Co., Inc., Novartis AG, Pfizer Inc., F. Hoffmann-La Roche, Schering-Plough, Synarc Inc., as well as nonprofit partners, the Alzheimer’s Association and Alzheimer’s Drug Discovery Foundation, with participation from the U.S. Food and Drug Administration. Private sector contributions to the ADNI are facilitated by the Foundation for the National Institutes of Health (www.fnih.org). The grantee organization is the Northern California Institute for Research and Education, and the study was coordinated by the Alzheimer’s Disease Cooperative Study at the University of California, San Diego. ADNI data are disseminated by the Laboratory for NeuroImaging at the University of California, Los Angeles. This research was also supported by NIH grants P30AG010129, K01 AG030514 and the Dana Foundation.

The OASIS data used in the preparation of this manuscript were obtained from the OASIS project funded by grants P50 AG05681, P01 AG03991, R01 AG021910, P50 MH071616, U24 RR021382, R01 MH56584. See http://www.oasis-brains.org/ for more details.

The AIBL data used in the preparation of this manuscript were obtained from the AIBL study of ageing funded by the Common-wealth Scientific Industrial Research Organization (CSIRO; a publicly funded government research organization), Science Industry Endowment Fund, National Health and Medical Research Council of Australia (project grant 1011689), Alzheimer’s Association, Alzheimer’s Drug Discovery Foundation, and an anonymous foundation. See www.aibl.csiro.au for further details.

The ICBM data used in the preparation of this manuscript were supported by Human Brain Project grant PO1MHO52176-11 (ICBM, P.I. Dr John Mazziotta) and Canadian Institutes of Health Research grant MOP-34996.

The IXI data used in the preparation of this manuscript were supported by the U.K. Engineering and Physical Sciences Research Council (EPSRC) GR/S21533/02 - http://www.brain-development.org/.

The ABIDE data used in the preparation of this manuscript were supported by ABIDE funding resources listed at http://fcon_1000.projects.nitrc.org/indi/abide/. ABIDE primary support for the work by Adriana Di Martino was provided by the NIMH (K23MH087770) and the Leon Levy Foundation. Primary support for the work by Michael P. Milham and the INDI team was provided by gifts from Joseph P. Healy and the Stavros Niarchos Foundation to the Child Mind Institute, as well as by an NIMH award to MPM (R03MH096321). http://fcon_1000.projects.nitrc.org/indi/abide/

This manuscript reflects the views of the authors and may not reflect the opinions or views of the database providers.

